# Cannabidiol pretreatment attenuates locomotor alterations and cytokine production in an autoimmune hepatitis model

**DOI:** 10.1101/2021.04.19.440455

**Authors:** Konstantinos Mesiakaris, Korina Atsopardi, Marigoula Margarity, Konstantinos Poulas

## Abstract

Cannabidiol (CBD) is a major active component of the Cannabis plant (*Cannabis Sativa L*.), which unlike tetrahydrocannabinol (THC) is devoid of euphoria-inducing properties. Broadly, CBD has demonstrated anxiolytic-like, anti-inflammatory and immunomodulatory effects. Concanavalin A (ConA) is a lectin found in the jack bean (*Canavalia ensiformis*) and it has been associated with a variety of toxicological effects (upon them mitogenic, cytotoxic and hepatotoxic). Intravenous administration of ConA is widely used for the induction of a model to study autoimmune hepatitis (AIH) in mice and the injury is mainly driven by activation and uptake of T-cells in liver. The aim of the present study was to investigate the effects of CBD administration (20 mg/kg), on adult mice, on locomotor activity and inflammatory markers, upon induction of AIH by ConA administration (20 mg/kg) on Balb/C mice. Inflammatory analysis was assessed by determining the IL-2, IL-4, IL-10 and INF-γ levels on plasma and sickness-like behavior assessed with open-field test. The results indicate that CBD pretreatment ameliorates impaired locomotor activity. IL-2, IL-4 and INF-γ levels on plasma were increased after ConA intoxication (inflammation index) and were reduced when mice were pre-treated with CBD. The detected IL-10 levels were increased when CBD pretreated, suggesting a protective anti-inflammatory effect.

## 1. Introduction

The biological activity of cannabis is due to a class of compounds called cannabinoids. Cannabidiol (CBD) is one of the 110 cannabinoids that were found in cannabis plant and accounts for up to 40% of the plant extract [1]. Despite the plethora of substances in hemp, the importance and the interest among cannabis products is focused on CBD content, due to its safety profile, the lack of intoxicating effects and its pharmacological properties, among them antioxidant, anti-inflammatory [2–4] and anxiolytic [5]. Its pharmacological effects are tested in various experimental models of diseases, among them inflammatory, autoimmune and neurodegenerative diseases and epilepsy [6]. One possible mechanism of its anti-inflammatory action is the deregulation of cytokine production by immune cells and the disruption of the physiologically regulated immune response. Endocannabinoids, metabolic enzymes, and their receptors have been identified in immune cells, monocytes, macrophages, basophils, lymphocytes and dendritic cells and they regulate immune function in an autocrine and paracrine way [7].

Inflammatory liver damage can be attributed to viral infections, autoimmune pathologies or drugs exposure and is the result of non-specific or even specific immune response [8]. Activation of T-lymphocytes and macrophages and cytokines production, are critical factors in the development of hepatitis and the disease progression is associated with pro-inflammatory cytokines secretion, such as IL-2, IL-6, IL-1β and TNF-α [9]. Autoimmune Hepatitis (AIH) is a chronic inflammatory liver disease [10,11] and its prevalence ranges from 50-200/millions cases and represents 5.9% of total liver transplants in the USA [12]. In this chronic inflammatory autoimmune condition, the hepatic cell destruction etiology remains unclear. The imbalance of immune system cells may be due to a combination of environmental and genetic factors, such as prescription drugs and infections. Several cytokines have been associated with its pathogenesis and severity [13,14], such as IL-2, IL-6, IL-1 β and TNF-α [9]. The immune response to autoantigens is not normally triggered and due to immune tolerance, autoantigens can be recognized by naive CD4-positive T cells [14]. Upon autoantigen recognition, naive T cells are activated and the immune reaction is initiated with differentiation into Th1, Th2, or Th17 cells, depending on the immunological microenvironment and the nature of the antigen epitope and these T cells trigger immune cascades [14]. Current standard treatment with azathioprine and prednisolone causes remission in most of the patients [11]. However, for those patients who do not respond to standard treatment or do not tolerate these drugs, few alternatives can be used and their efficacy may be limited [11].

The anti-inflammatory profile of CBD, as well as the research on animal models of inflammatory diseases, have led to this study of CBD administration in a model of autoimmune hepatitis. Concanavalin A (ConA) is a lectin found in the jack bean (*Canavalia ensiformis*) with a variety of toxicological effects and ConA murine model is a well-established model, which resembles the pathophysiology of immunologically mediated liver disorders such as AIH [15]. Upon intoxication, ConA decreases locomotor activity, which is translated as sickness behavior, a coordinated set of adaptive behavioral changes triggered by an activation of the peripheral innate immune system and the production of pro-inflammatory cytokines [16–18]. Sickness behavior is associated with the behavioral changes, observed both in humans and laboratory animals, that accompany the response to infection and is expressed by symptomatology of depressive-like behavior, anhedonia, fatigue, psychomotor slowing, decreased appetite, sleep alterations and increased sensitivity to pain [19,20]. In this behavior, cytokines are responsible for this symptomatology observed in response to inflammation (eg, anorexia, decreased motor activity [27]. The main cells that are activated after concanavalin A administration are the CD4 activated T-cells and the natural killer cells (NK), and in conjunction with macrophages and eosinophils, they induce hepatic cells death directly or indirectly through the production of high levels of inflammatory cytokines [21–23]. Controlling the secretion of inflammatory and anti-inflammatory cytokines, could be an important step in managing autoimmune hepatitis. To further support the notion that pro-inflammatory cytokines are the key mediators of sickness behavior, administration of cytokine modifiers, such as cytokine antagonists, can result in changes in sickness behavior [20]. Administration of cytokine modifiers, such as cytokine antagonists, can lead to changes in sickness behavior, a fact that supports that pro-inflammatory cytokines may be the key mediators of sickness behavior.

CBD research indicates its anti-inflammatory potential in various experimental models, but there is not information describing the effects of CBD on the model of AIH of Concanavalin A. This study aims to examine the potential changes in cytokine production and sickness behavior, after CBD administration, in this model and the possibility of CBD as a therapeutic tool in managing autoimmune hepatic disorders.

## 2. Materials and methods

### 2.1. Reagents

Concanavalin A and the other necessary reagents were supplied by Sigma-Aldrich. CBD supplied by Cayman chemical (CAS No. 13956-29-1). The Mouse Th1/Th2 Uncoated ELISA kit was acquired from ThermoFisher Scientific (Catalog Number 88-7711-44).

### 2.2. Animals

Male Balb-c mice [22-30 g body weight (BW)], 3–4-month-old, were housed in standard laboratory polyacrylic cages (3–5 mice per cage), with free access to water and food. Mice were kept in a room under controlled temperature (22 ± 1 °C), relative humidity of 50–60%, and a controlled light–dark cycle (12 h/12 h). The handling of animals and the study protocol were in accordance with Greek Presidential decree 86/2020, for adaptation of Greek legislation to directive 2010/63/EU of the European parliament and of the council of 22 September 2010 on the protection of animals used for scientific purposes. All efforts were made to minimize the number of animals and for the optimization of living conditions and optimal manipulations were held to eliminate stress.

### 2.3. Study design

Cannabidiol (CBD) administered orally with gavage, once daily for a period of 5 days in dose 20 mg/kg. Concanavalin A (ConA) was dissolved in a vehicle solution (saline, 0.9% NaCl) making a stock solution of 10 mg/ml and was administered intravenously (i.v.) immediately after dissolution, at a dose of 20 mg/kg. The number of animals used in each group was five (n=5). Mice were randomly separated into five groups (5 different mice/group) and treated as follows: In Control CBD group, mice received water orally with gavage for 5 consecutive days (once daily). In CBD group, mice received CBD orally (20 mg/kg body weight/day) with gavage for 5 consecutive days (once daily). In Control ConA group, mice received once Intravenous (IV) injection of saline (0.9% NaCl). In ConA group, mice received once Intravenous (IV) injection of Concanavalin A (20 mg/kg). In ConA + CBD group, mice received orally CBD (20 mg/kg body weight/day) with gavage for 5 consecutive days (once) and 2 h after the last CBD dose, mice received once Intravenous (IV) injection of Concanavalin A (20 mg/kg).

Behavioral tests were performed 3h after the last administration of CBD (5th day) in CBD groups (Control CBD, CBD and ConA + CBD) and 2 hours after the IV doses in ConA groups (Control ConA and ConA). Animals were then led to euthanasia by cervical dislocation. Blood collected immediately with cardiac puncture and plasma collected with centrifugation at 1000 g for 10 minutes at 4 °C and stored at – 80 °C for further analysis.

### 2.4. Locomotor Activity analysis

Locomotor Activity was assessed using the open-field apparatus. Open field behavior is applied as an index of locomotor activity in animal models of hepatic disorders which result in impaired activity [24–26]. On the day of the behavioral test, mice were initially kept in their cages for 1 h in a slightly illuminated and sound-proof room to adapt to the new environment. At the beginning of the test, each animal was carefully placed in the central area of the device, and its locomotor activity was recorded for 10 min. Every time that a mouse was entering the center of the apparatus was counted as an entrance and the recordings were assessed as an index of locomotor activity.

### 2.6. Statistical analysis

Statistical analysis performed by the IBM SPSS statistical package V24.0, and the graphs were constructed in Microsoft Excel 365. One-way analysis of variance (ANOVA) followed by tukey post hoc comparisons test was used for statistical analysis in behavioral (anxiety-like behavior) and biochemical assays. Data are presented as mean ± S.E.M. Probability values of less than p < 0.05 were considered statistically significant, as the statistical significance level has been set the 95%.

## 4. Results

### 4.1. Effect of cannabidiol on behavioral parameters

CBD effect on locomotor activity assessed either with or without concanavalin intoxication, to estimate CBD effect in both healthy and AIH induced groups. As shown at fig.1, cannabidiol administration increases locomotor activity, with the increase of the number of entries to the center compared to the control. Moreover, concanavalin A administration reduces locomotor activity. Changes in locomotor activity after ConA administration may be associated with a decrease in Tyrosine Hydroxylase (TH) levels in brain, through the release of cytokines such as IFN-γ and TNFα TNFα [16]. Cannabidiol pre-administration at AIH mice increases the number of entries, and this means an increase of mobility at levels comparable to control ConA group.

**Figure 1.**
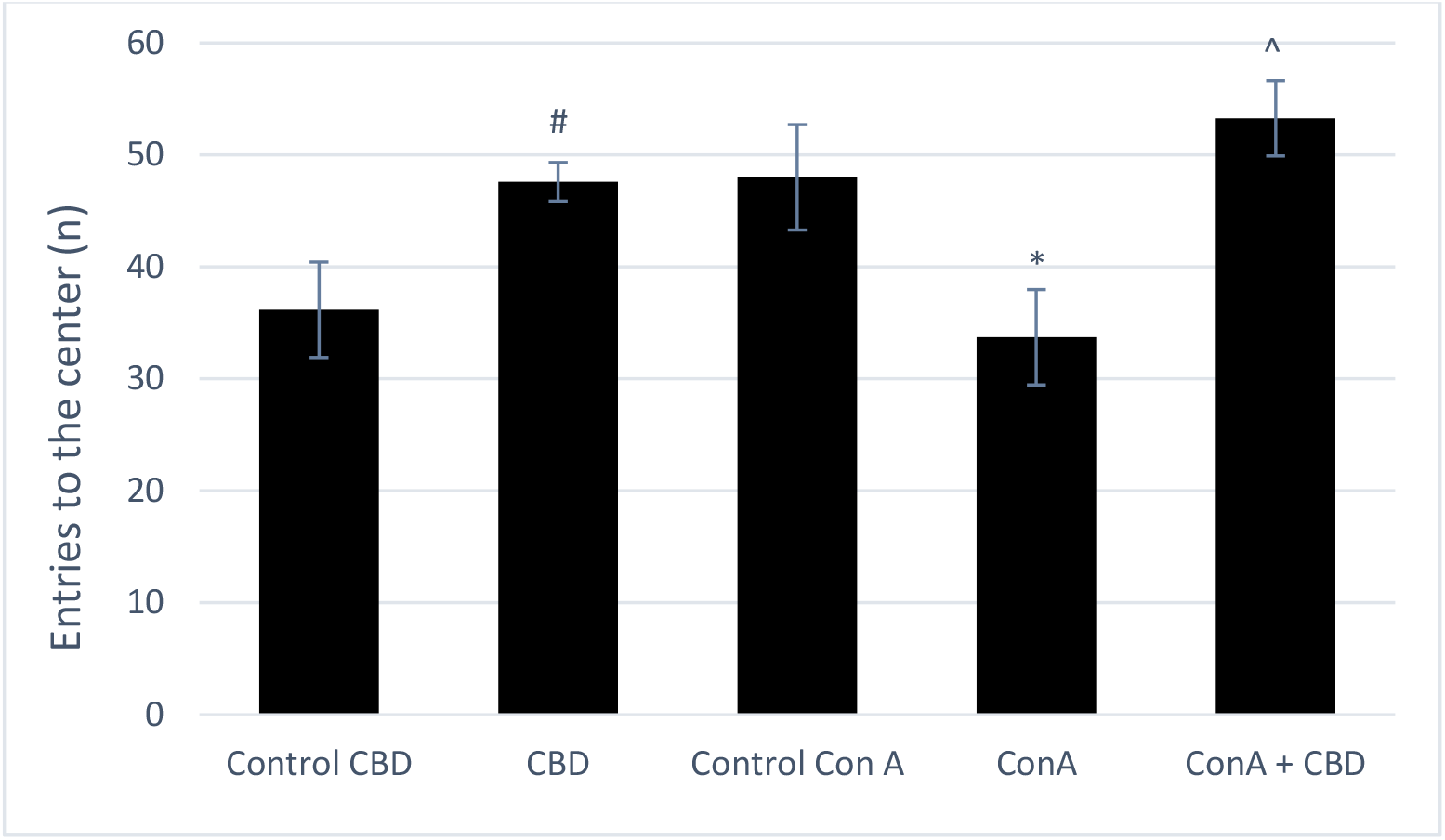
CBD administration (20 mg / kg) effect on locomotor activity in healthy subjects and AIH model. The bars correspond to the mean value ± standard error, on 5 animals in each experimental group (n = 5). *: p <0.05 (T-student test) compared to control CBD group, #: p <0.05 (Anova) compared to Control ConA group, ^: p <0.05 (Anova) compared to ConA group

### 4.2. Effect of cannabidiol administration on cytokines levels in AIH model

The results of cannabidiol administration on cytokine production in the AIH model are presented in fig. 2. AIH induction increases interferon-γ levels in AIH model, ConA group (7230.04 pg/ml). and Although CBD pretreatment in AIH model does not change these increased levels, ConA+CBD group (6217.95 pg/ml). IL-2 levels are detected in all experimental groups, control CBD (213.89 pg/ml), CBD (254.25 pg/ml) and Control ConA (6.18 pg/ml) and presents statistically increased levels in ConA group (5914.56 pg/ml) and statistically decreased levels in ConA+CBD (3107.22 pg/ml) in compare to ConA group. IL-4 levels are also increased after AIH induction at ConA group (1796.50 pg/ml). CBD pretreatment results at significantly lower levels of IL-4 in ConA+CBD group (423.34 pg/ml). Low IL-10 level detected in AIH model in ConA group (304.00 pg/ml) while cannabidiol pretreatment results in significantly increased level in ConA+CBD group (620.50 pg/ml).

**Figure 2.**
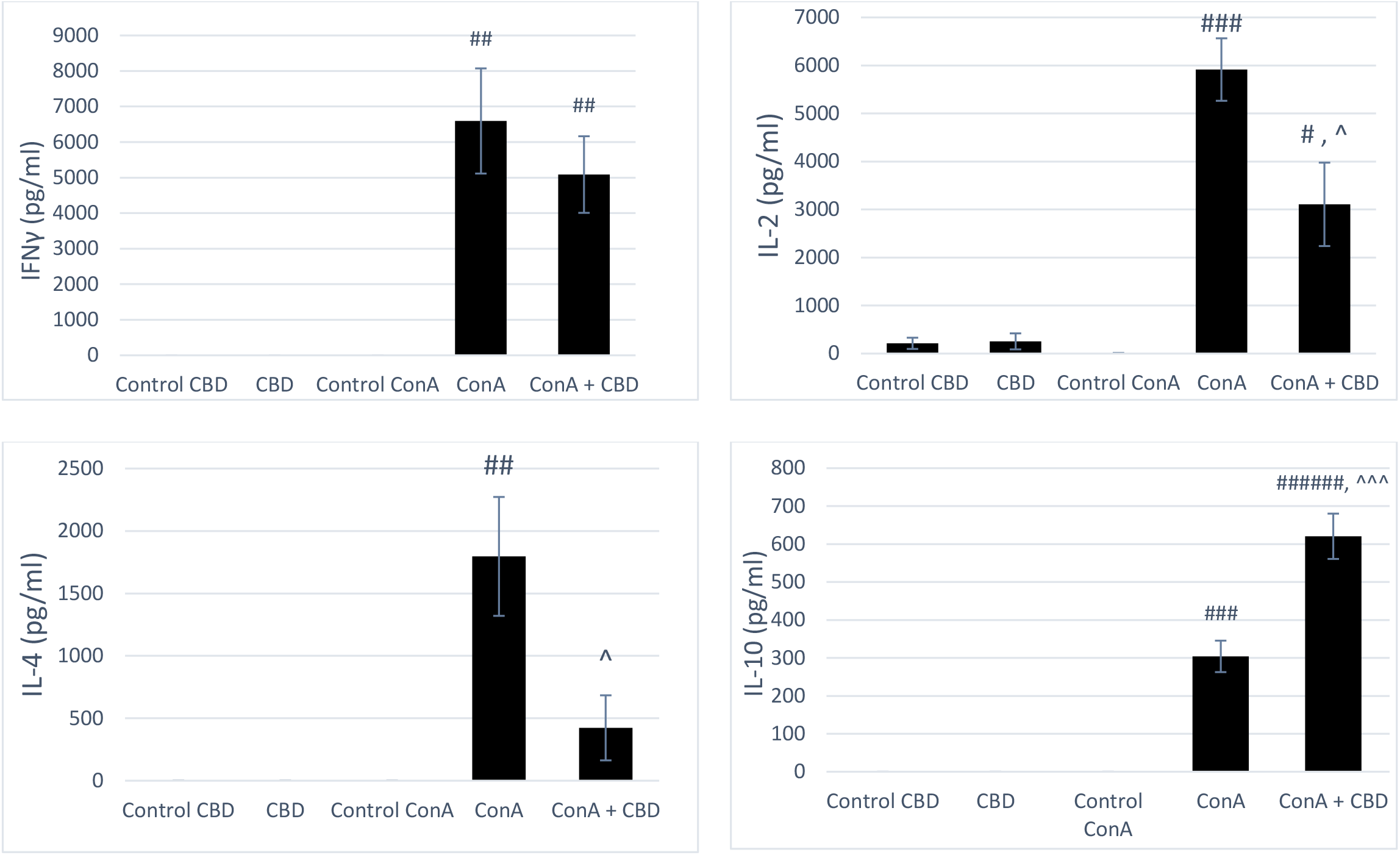
CBD administration (20 mg / kg) effect on cytokine production in healthy subjects and AIH model. The bars correspond to the mean value ± standard error, on 5 animals in each experimental group (n = 5). #: p <0.05, ##: p<0.01, ###: p<0.001, ######: p<0.0000001 compared to Control ConA group (One-Way Anova Analysis), ^: p <0.05, ^^^: p<0.001 compared to ConA group (One-Way Anova Analysis).

## 5. Discussion

The present study examined the effect of cannabidiol on cytokine levels, as well as its effect on behavioral markers in an induced model of autoimmune hepatitis. TNF-α and IFN-γ are key factors in ConA-induced liver damage [22,28]. Following ConA administration, IL-2, IL-4 and IFNγ levels rise sharply [29], as it is observed at the results above. IL-10 has been classified as an anti-inflammatory cytokine [8,30–32] and at this model it prevents the development of liver damage as it was expected [33,34]. That agrees with the results as when CBD is pretreated, IL-10 levels are increased and at the same time IL-2 and IL-4 levels are decreased. It is previously described that IL-2 reduction in the ConA model, after administration of a protective agent, is explained as prevention of ConA-induced hepatitis [9]. IL-4 is thought to be a pro-inflammatory cytokine, which recruits more inflammatory cells, such as eosinophils and macrophages, involved in inflammatory responses [35–37]. Thus, the reduction observed in IL-4 levels, when cannabidiol is administered as pretreatment, indicates a possible anti-inflammatory protective effect of CBD. It is known that peripheral inflammatory conditions, chronic or acute, may alter the mood and motility of the subjects [38]. ConA administration results in impaired locomotor activity, as well as the manifestation of a behavioral alterations [27]. According to the results presented above, measuring the locomotor activity results in increase of it when CBD is administrated. This suggests that CBD administration improves sickness behavior caused by ConA in the AIH model.

## 6. Conclusions

Excessive responses can lead to uncontrolled tissue damage, so there must be a mechanism for balancing them. This study suggests that CBD administration attenuates AIH sickness behavior with a downregulation of IL-2 and IL-4, with an upregulation of IL-10. Controlling the secretion of inflammatory cytokines, as well as anti-inflammatory cytokines, such as IL-10, could be an important step in managing autoimmune hepatitis and autoimmune hepatic disorders.

## Acknowledgments

None

